# Impact of protein conformational diversity on AlphaFold predictions

**DOI:** 10.1101/2021.10.27.466189

**Authors:** Tadeo Saldaño, Nahuel Escobedo, Julia Marchetti, Diego Javier Zea, Juan Mac Donagh, Ana Julia Velez Rueda, Eduardo Gonik, Agustina García Melani, Julieta Novomisky Nechcoff, Martín N. Salas, Tomás Peters, Nicolás Demitroff, Sebastian Fernandez Alberti, Nicolas Palopoli, Maria Silvina Fornasari, Gustavo Parisi

**Author notes:** corresponding author: Gustavo Parisi. authors contributed equally (Tadeo Saldaño, Nahuel Escobedo).

## Abstract

After the outstanding breakthrough of AlphaFold in predicting protein 3D models, new questions appeared and remain unanswered. The ensemble nature of proteins, for example, challenges the structural prediction methods because the models should represent a set of conformers instead of single structures. The evolutionary and structural features captured by effective deep learning techniques may unveil the information to generate several diverse conformations from a single sequence. Here we address the performance of AlphaFold2 predictions under this ensemble paradigm. Using a curated collection of apo-holo conformations, we found that AlphaFold2 predicts the holo form of a protein in 70% of the cases, being unable to reproduce the observed conformational diversity with an equivalent error than in the estimation of a single conformation. More importantly, we found that AlphaFold2’s performance worsens with the increasing conformational diversity of the studied protein. This impairment is related to the heterogeneity in the degree of conformational diversity found between different members of the homologous family of the protein under study. Finally, we found that main-chain flexibility associated with apo-holo pairs of conformers negatively correlates with the predicted local model quality score plDDT, indicating that plDDT values in a single 3D model could be used to infer local conformational changes linked to ligand binding transitions.

## Introduction

Attempts to predict protein structure from their sequences started in the early sixties with Anfinsen’s experiment, which showed that the structure of a protein is encoded in its amino-acid sequence [1]. After decades of extensive experimentation and efforts, the practical demonstration of Anfinsen’s motto came from deep learning techniques taking advantage of evolutionary information. In the last year, the computational tool AlphaFold2 [2] developed by DeepMind, reached an impressive performance in predicting protein structures with an accuracy similar to experimental techniques [3,4]. AlphaFold2 is based on a novel neural network architecture that attends over evolutionary information, codified in an MSA, to create a novel representation of the sequence and the relative distances between residues. Those representations are further improved using an end-to-end approach to generate structure models with iterative structural refinement. The output of AlphaFold2 is a highly accurate set of structural models with accompanying residue-specific estimates of modeling reliability.

This outstanding achievement is not only conceptual, in the sense of the advancement of novel deep learning techniques and protein science, but also practical. It provides the scientific community with a method for fast, reliable, and cheap determination of structural models that can be applied at a large scale. Recently, DeepMind and EMBL-EBI have jointly released the database of AlphaFold2 predictions for the whole human proteome [5] and other key organisms (https://alphafold.ebi.ac.uk/). Furthermore, an easy-to-use and fast version of the AlphaFold2 pipeline was introduced by modifying the time-consuming step of multiple sequence alignments generation with almost identical results [6]. These exceptional endeavors will soon contribute to filling the gap between proteomes and structuromes, triggering the blooming of almost every related biology field involving both wet-lab practices and computational-based approaches.

The AlphaFold2 neural network is trained using structures derived from crystallization and X-ray diffraction experiments. It is thus expected that the 3D models obtained will reproduce “regular” PDB structures [2]. How much do regular PDB structures resemble the native state of proteins? It is widely accepted that protein function relies on a conformational ensemble describing the native state of proteins [7–10] that is not entirely captured in the PDB [11]. Structural differences between conformers promote ligand binding [12], transport [13], or catalysis [14,15]. These differences are also relevant for signal transduction [16] and define metabolic regulation by mechanisms like cooperativity and allosterism [10,17–19]. Conformers in the native ensemble could be identical in their backbones but differ just in the conformations of some residues, defining open and close transitions of tunnels [20,21] and/or volume variations in their cavities [22,23]. Increasing differences involve backbone movements comprising loops, secondary structural elements rearrangements, and relative domains movements [24–26]. Extreme cases of conformational diversity are represented by intrinsically disordered proteins which lack tertiary structure and form complex ensembles with high ratios of interchange between conformers [27].

Given the ensemble nature of proteins we explore the impact of conformational diversity in the AlphaFold2 performance prediction. Firstly, we relied on a hand-curated set of proteins with different extensions of experimentally estimated conformational diversity, defined by an apo conformer and the corresponding holo form bound to a biologically relevant ligand. Using this dataset, we studied if AlphaFold2 can reproduce different conformers among their resulting top-scoring models. We also explored how AlphaFold2’s performance is affected by the degree of conformational diversity of the protein under study. Additionally, as AlphaFold2’s predictions heavily rely on evolutionary information, we used families of homologous proteins with different extensions of conformational diversity among its members to test whether this heterogeneity affects prediction.

## Results

### Description of the dataset

We selected 86 proteins (Supplementary Table 1) with different degrees of conformational diversity expressed as the range of pairwise global Cα-RMSD between their conformers in the PDB (Figure 1). All the pairs of conformers for each protein are apo-holo pairs selected from the CoDNaS database [28] and bibliography. Hand-curation for each protein confirmed that structural deformations were associated with a given biological process based on experimental evidence. This step is essential to ensure that conformational diversity is not associated with artifacts, misalignments, missing regions, or the presence of flexible ends. When more than two conformers are known, we have selected the apo-holo pair showing the maximum Cα-RMSD (maxRMSD). Other considerations were absence of disorder, PDB resolution, absence of mutations, and sequence differences. We previously observed that when conformational diversity is derived from experimentally-based conformers, different extensions of RMSD are obtained between them depending on the structure determination method [20]. Here we considered a continuum of protein flexibility measured as the RMSD between apo and holo forms as shown in Figure 1.

**Figure 1:**
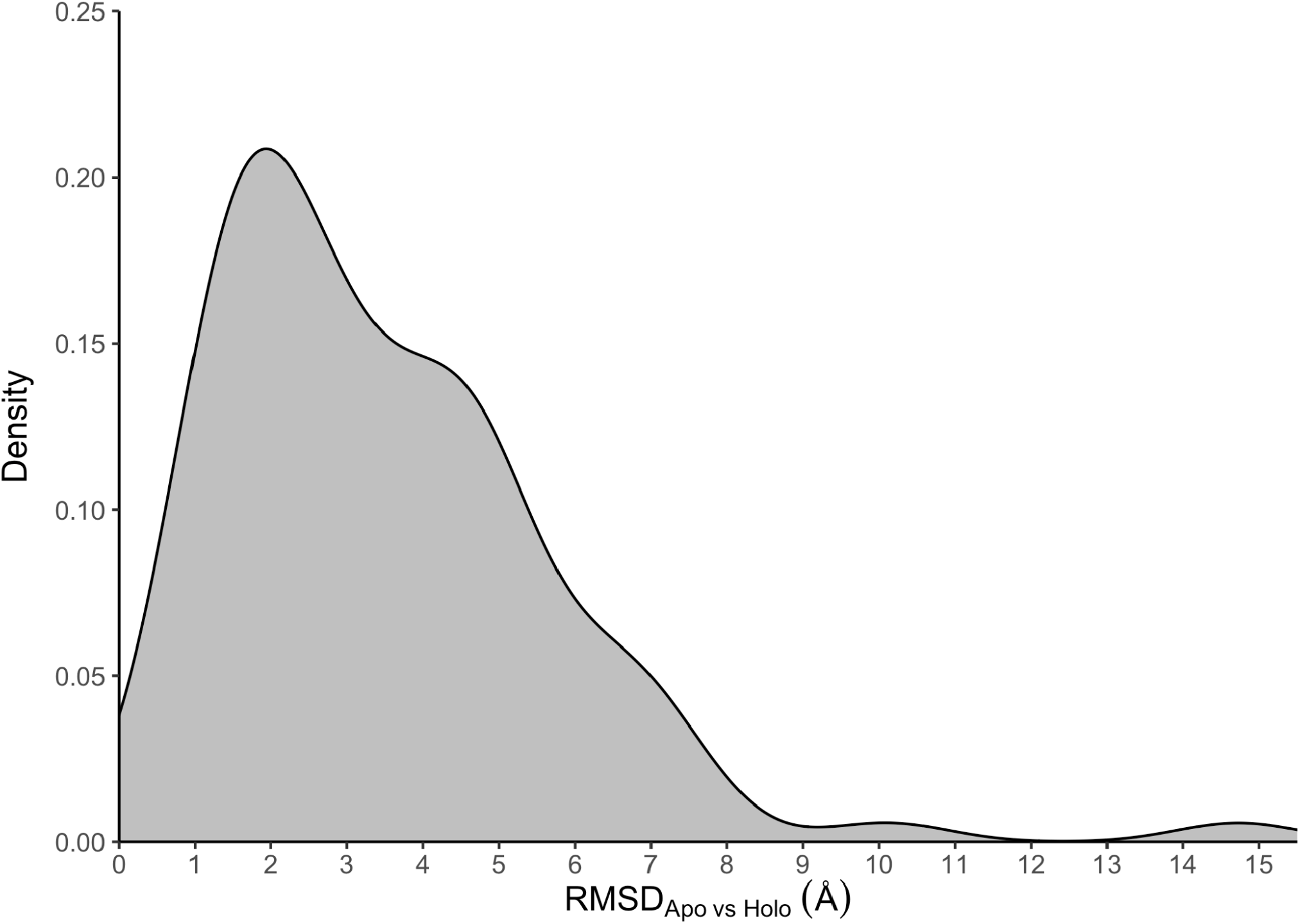
Distribution of RMSD between apo-holo pairs. The average of the distribution is 3.5Å.

### AlphaFold2 does not reproduce conformational diversity

We have predicted the structure of each protein in the dataset using ColabFold (AlphaFold2.ipynb - Colaboratory) running AlphaFold2 with MMSeq2 [6] without the use of templates and with the option to obtain relaxed models with Amber force fields [29], gathering the first five top models according to plDDT (predicted local Distance Difference Test) scores [30]. Supplementary Figure 1 shows the distribution of the plDDT scores for all the models. We found that 90% of the models scored higher than 86.9, reaching 90.0 if only the best model for each protein is considered, evidencing the good quality of the models obtained.

All AlphaFold2 models (five per protein) were structurally aligned to the experimentally resolved apo and holo conformations for each protein in the dataset, and the RMSD value was calculated for each alignment. Figure 2A shows the relationships of RMSD values against the apo and holo forms for all the obtained AlphaFold2 models, while Figure 2B is limited to the best model for each protein, according to the plDDT.

**Figure 2:**
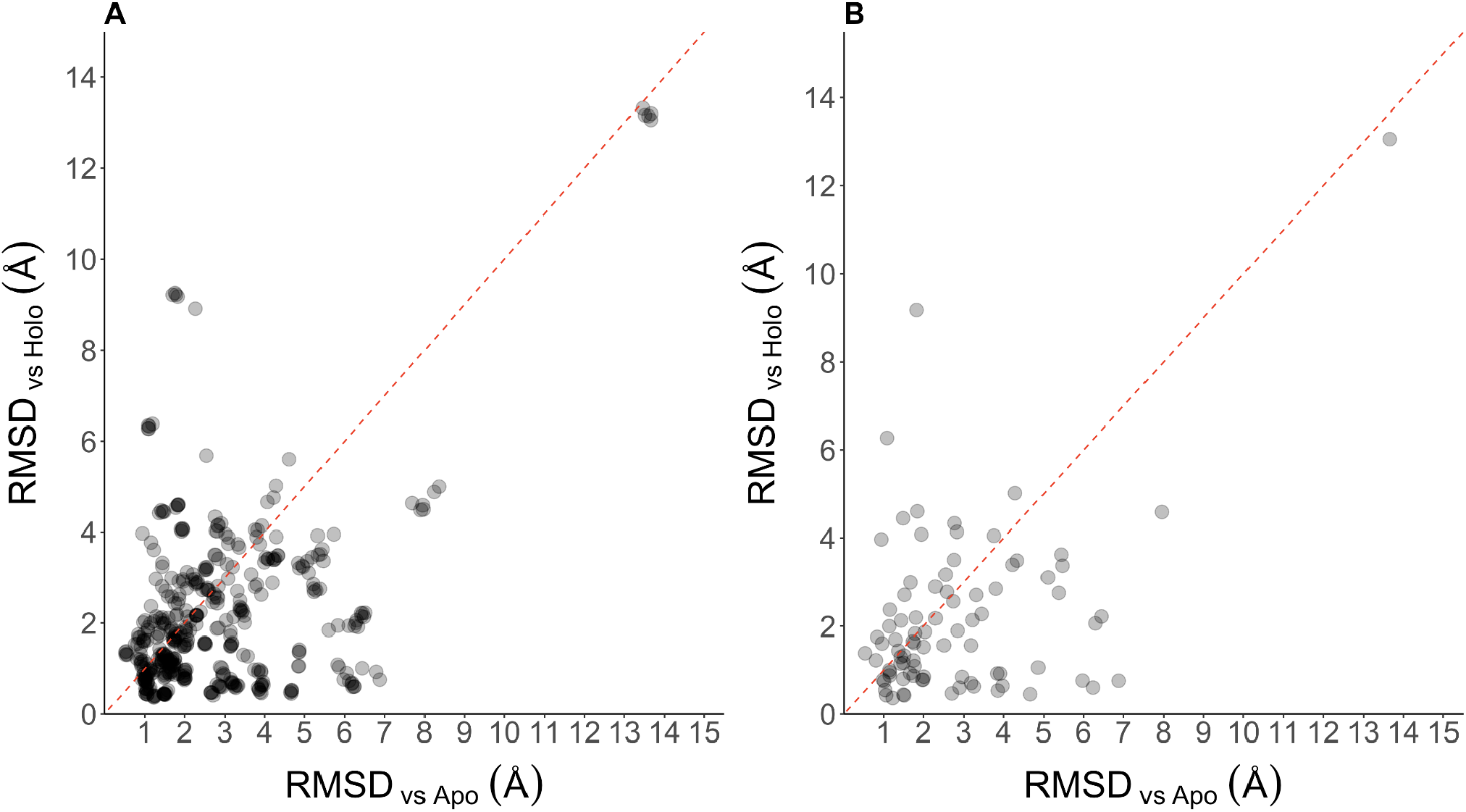
Correlation between RMSD values derived from the alignment of each AlphaFold2 model to the apo (x axis) and holo (y axis) conformations of each protein. **Panel A** shows the distribution of RMSD for all the models, while **panel B** is limited only to the best models according to pIDDT scores.

According to these RMSDs, we found that 67% of the proteins show a model with the lowest RMSD to the holo form and 33% to the apo form. This tendency is maintained when only the highest scoring model (in terms of plDDT) is considered (69.8% and 30.2% for holo and apo form).

In Figure 3, we plot the distributions of the best RMSD obtained between the models with their apo and holo forms, discriminating among proteins that were modelled closer to the apo (left) or to the holo (right) forms. For proteins modelled closer to the apo form, the average of the lowest RMSDs between the models and the apo and holo forms is 1.94Å and 3.59Å, respectively (Figure 3, left). On the contrary, for proteins modelled closer to the holo form, the average of the lowest RMSDs between the models and the holo form is 1.79Å and climbs to 3.36Å against the apo form (Figure 3, right). We conclude that most of the proteins are modelled with a bias towards a given conformer. It is then impossible to estimate the degree of conformational diversity captured in apo and holo pairs with the same precision that can be estimated for a single representative conformation of a given protein. As expected, the error in the estimation of the conformational diversity is highly correlated with the structural differences between the apo and the holo forms. The correlation between the conformational diversity of the protein with the best RMSD measured to the unfavored form (see Figure 3) has a 0.96 Pearson Correlation (p-value < 0.001) (See supplementary Figure 2).

**Figure 3:**
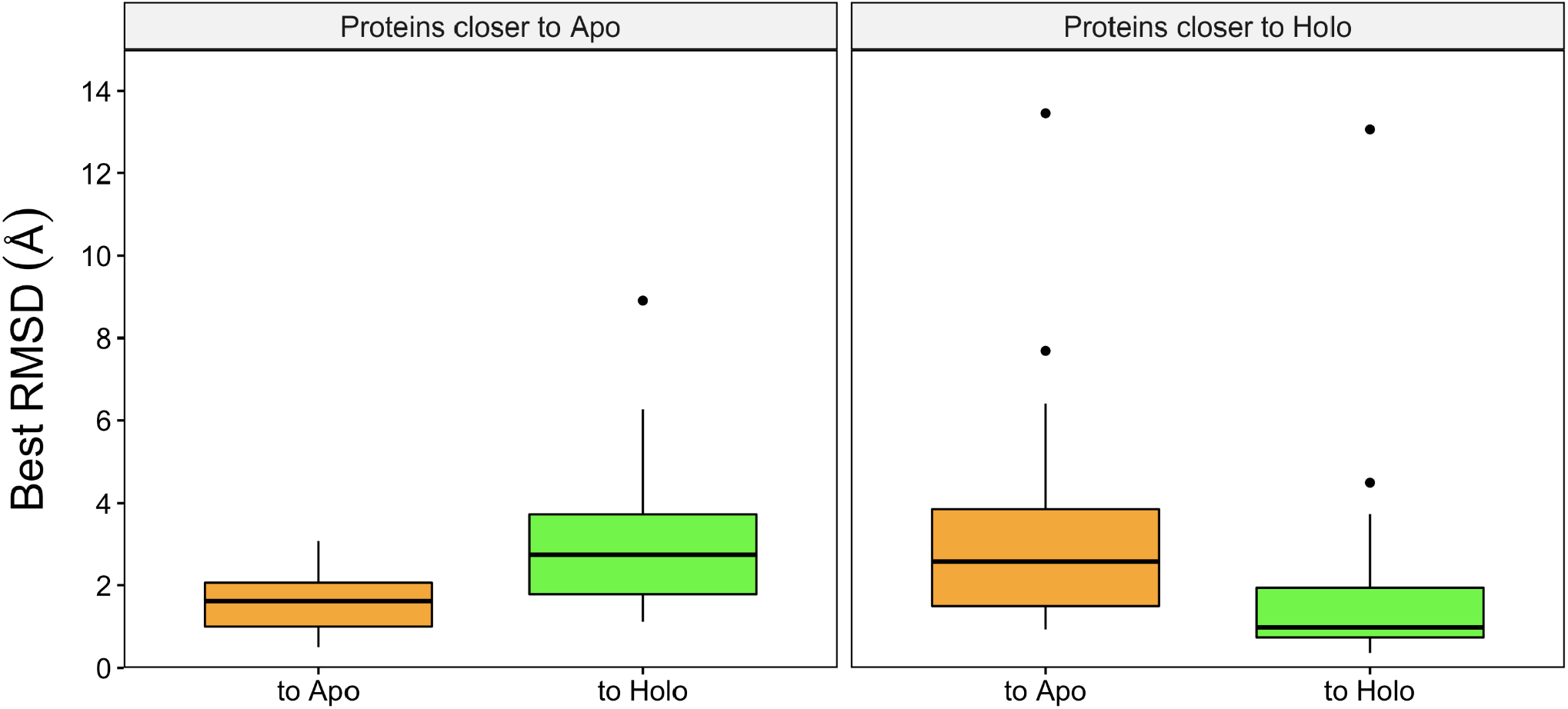
Distribution of best RMSD between apo and holo forms and estimated models. “To Apo” and “To Holo” are the average lowest RMSD towards apo and holo forms respectively. The different panels involve proteins with a model showing best RMSD among models towards apo form (left panel) and towards holo form (right panel).

The preference for a single conformer, apo or holo, is not associated with the AlphaFold2 predictive performance since the median of the pIDDT score distribution for models that resemble the holo and apo forms are 95,47 and 94.53, respectively (Wilcoxon p-value = 0.0315). When using the best models only, the corresponding medians are 96.33 and 95.96, respectively (Wilcoxon p-value = 0.494).

Figure 4 shows three examples that illustrate the model preference for the apo or the holo form, or its lack, taken from the results described above.

**Figure 4:**
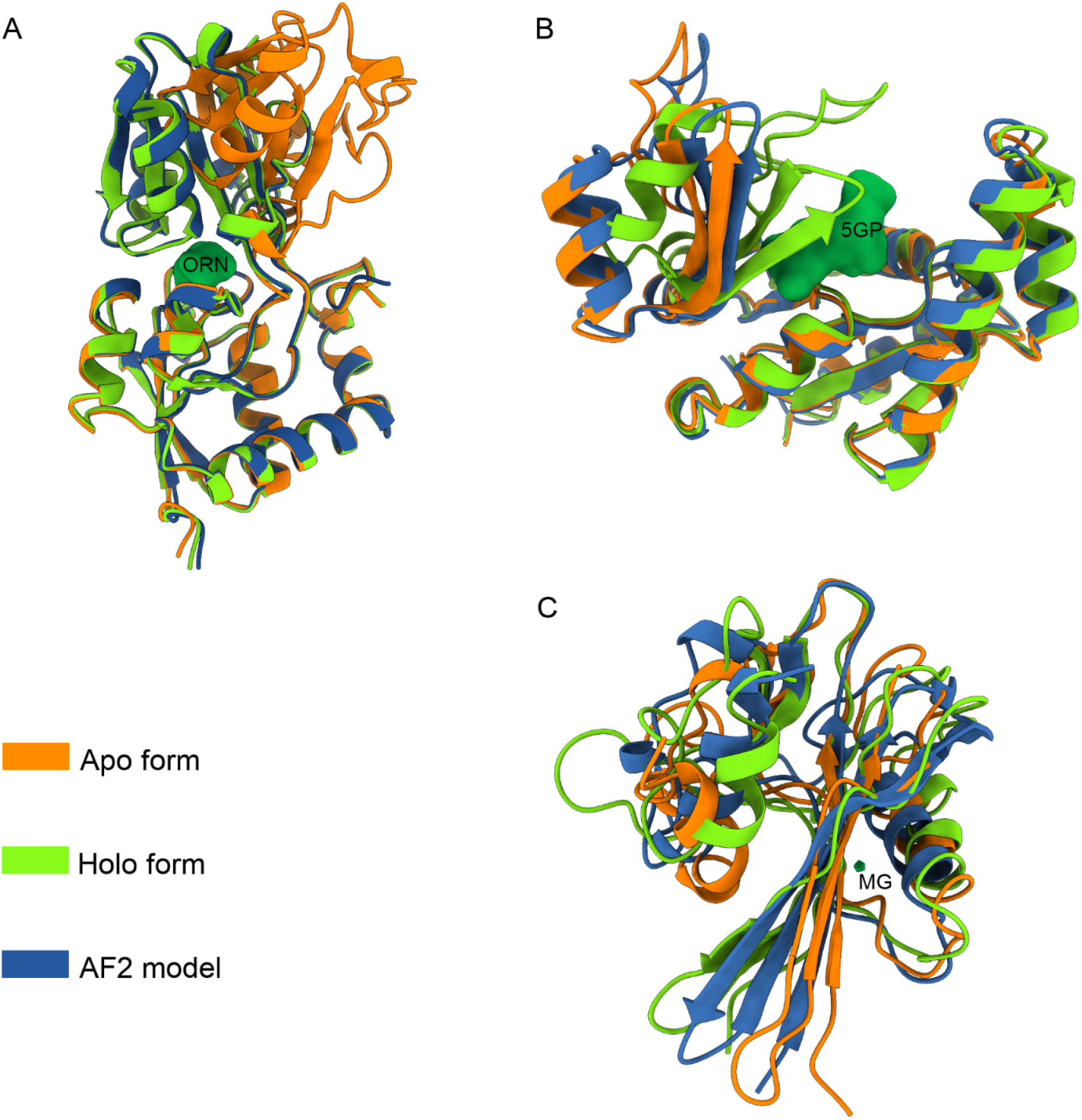
Three different situations of predictions respect experimental structure. **A**. The AlphaFold2 model closest to an experimental structure is closer to the holo (Cα-RMSD = 0.45Å, PDB ID = 1LAH_E) than the apo form (4.67Å, 2LAO_A). Apo and holo structures correspond to the multiple ligand specificity of the periplasmic lysine-, arginine-, ornithine-binding protein (LAO) [31]. **B**. Guanylate Kinase from yeast, where the best AlphaFold2 model showed better match with the apo (Cα-RMSD = 0.94Å, PDB ID = 1EX6_B) form than with the holo form (3.97Å, 1EX7_A) [32]. **C**. A case where AlphaFold2 model is different to both the apo (Cα-RMSD = 3.76Å, PDB ID = 1MUT-11_A) and holo (4.05Å, 1PUN-7_A) forms of a nucleoside triphosphate pyrophosphohydrolase from E. coli [33]. Proteins are shown as cartoons while biologically significant ligands are labelled and shown in surface representation.

### AlphaFold2 predictions worsen with increasing conformational diversity of the protein

Given that AlphaFold models better resemble the holo conformation of proteins, in this section, we studied how the conformational diversity of the protein, measured as the structural differences between apo and holo forms(RMSDapo_holo), affects performance predictions. We found that models are less predictable as RMSDapo-holo increases, measuring the prediction error as the lowest RMSD (lowestRMSD) of a model to the apo or holo forms (Figure 5A). We found that the predictive performance highly depends on the conformational diversity of the protein (0.69 Pearson correlation coefficient, p-value <0.001). This tendency is also observed when we studied the correlation between the global plDDT for the best model and the RMSDapo_holo (−0.53 Pearson correlation coefficient and p-value <0.001) (Figure 5B).

**Figure 5:**
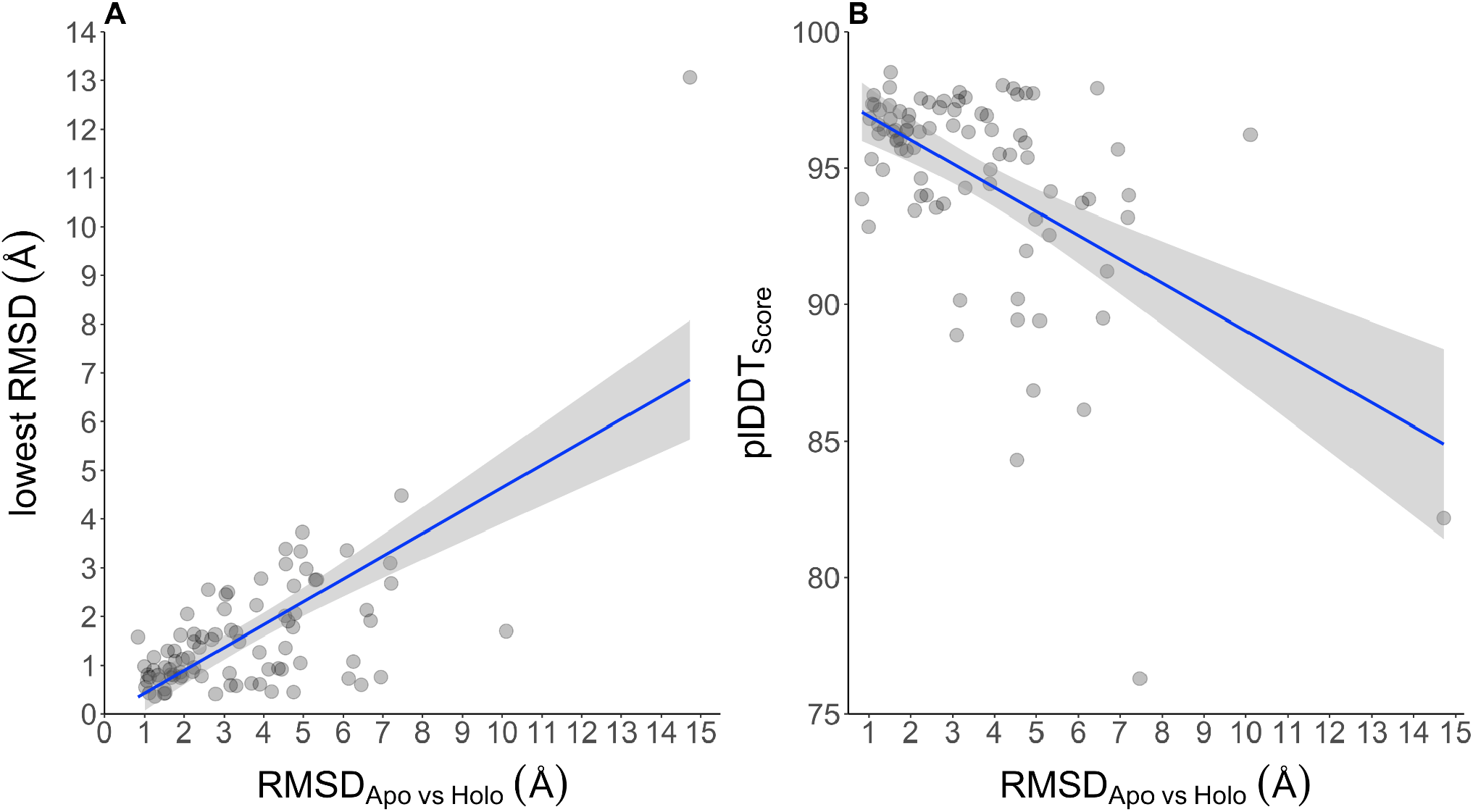
Quality prediction of AlphaFold2. **Panel A**: As conformational diversity increases between apo and holo forms, the lowest RMSD to any of the forms increases as well (Pearson correlation 0.69, p-value < 0.001). **Panel B**: Likewise, the global plDDT scores decrease with larger protein conformational diversity (panel b).

The model error shows no dependency on the sequence number of the input alignment (−0.18 Pearson correlation coefficient, p-value >0.05) and protein length (−0.25 Pearson correlation coefficient, p-value <0.05). To study how the error in the model depends on the type of protein movements between the apo and holo forms, we classified the pairs in our dataset in two broad categories [34]: according to the presence of domains and hinges movements using the DynDom software [35] and the presence of flexible loops in just one domain. We found that the error measured as the lowestRMSD is slightly dependent on the type of movement (Wilcoxon rank sum test W= 1128.5, p-value = 0.046) with a median of 0.93 and 1.6Å for the domain movements group and proteins with loop movements.

### Fuzzy evolutionary information could affect AlphaFold2 prediction

To explain the impairment observed in AlphaFold2 prediction capacity with increasing protein conformational diversity, we hypothesized that the evolutionary information in the input multiple sequence alignment could be fuzzy due to the conformational diversity heterogeneity in the protein family. Previously we have found that families with highly flexible proteins [20] heavily affect homology modeling due to a noisy relationship between sequence and structure divergence [36]. Moreover, we have also characterized that the inter-residue contacts predicted using coevolutionary methods are the consensus ones, independently of the structural variations among family members [37]. Taking into account the finding that protein dynamical behaviour is mostly not conserved in protein families [36], families that include highly flexible and rigid proteins could have confounding mixtures of sequence signatures.

To test this hypothesis, we explore families of homologous proteins with experimentally based conformational diversity. These families were obtained from the CoDNaS database using a sequence-based clustering with 40% sequence identity and 70% coverage. Each of the ∼29000 protein entries in CoDNaS has an associated maximum Cα-RMSD derived from comparing all the conformers belonging to a given protein. This maxRMSD is taken as the maximum conformational diversity the protein could have. Clusters were further classified into homogeneous or heterogeneous according to the range between the minRMSD and the maxRMSD for each protein in each family (range < 4Å for homogeneous families, or heterogeneous otherwise).

Mapping our 86 proteins to these clusters only retrieved 20 proteins distributed in 10 homogeneous and 10 heterogeneous clusters. For each of these 20 proteins we study the correlation between their corresponding error in AlphaFold2 predictions (estimated above as the lowest RMSD to any experimental conformer) and the dispersion of the conformational diversity in all proteins from the same family. We found that the heterogeneous clusters performed worse than the homogeneous ones (average lowestRMSD values for hetero and homogeneous families are 2.03Å and 1.31Å, respectively; Wilcoxon p-value <0.005).

To further test this hypothesis, we repeated the estimation with 175 chosen proteins, one for each of the most populated clusters described above (23.53 homologous proteins on average per cluster). For each of these 175 proteins, we ran AlphaFold2 using ColabFold and estimated the lowest RMSD, comparing the top obtained models with the corresponding structure of the protein. We observed the same trend as shown in Figure 6A (average lowest RMSD of 1.94Å and 1.54Å for hetero and homogeneous families, respectively; Wilcoxon p-value < 0.005).

**Figure 6:**
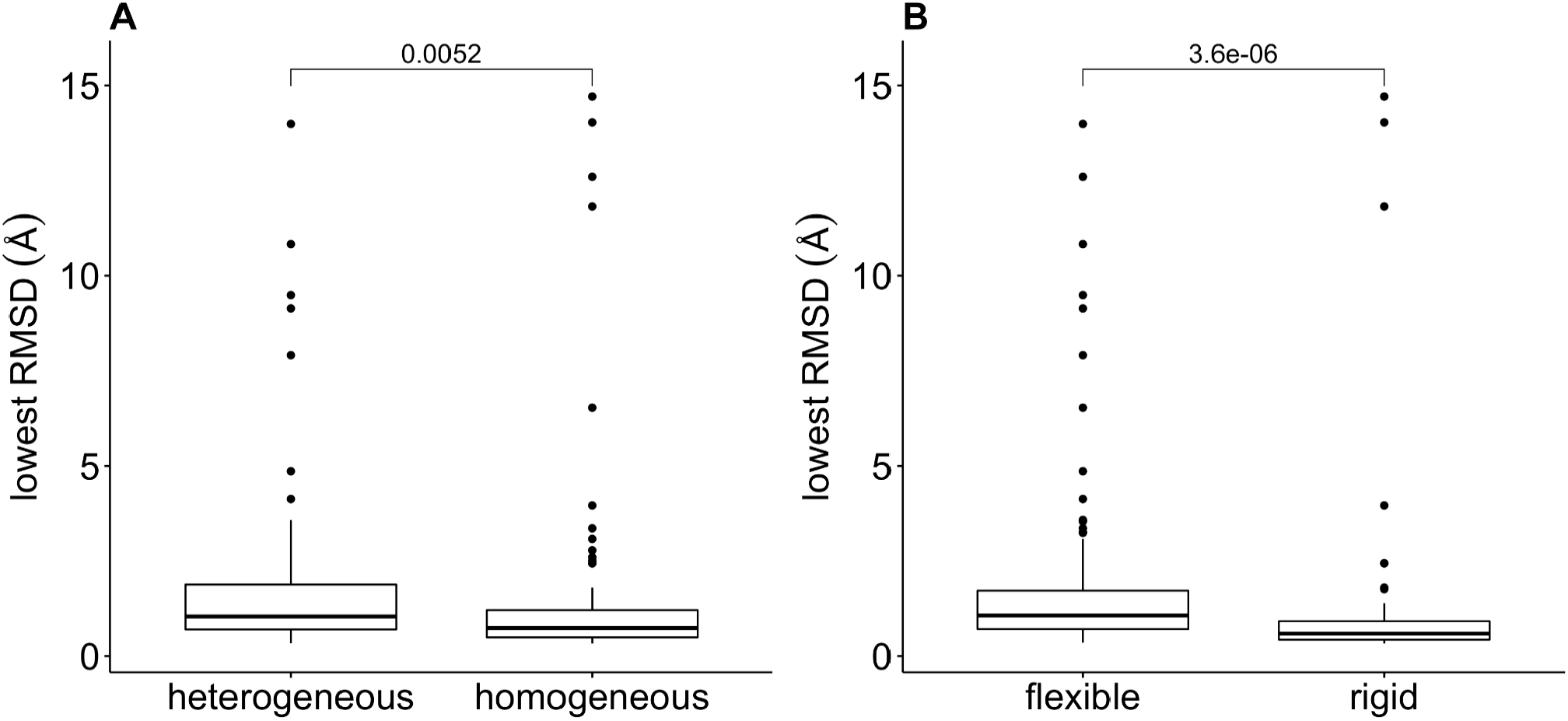
Model error estimated as the lowest RMSD to the apo and holo forms as a function of the evaluation of the distribution of the conformational diversity in 175 homologous families. Panel A contains families classified as heterogeneous and homogeneous using the range of the proteins in each family, while in Panel B they were classified as flexible and rigid.

To further explore if the model estimation is affected by the flexibility of each family, we further classified the clusters in “flexible” and “rigid”, with a flexible cluster defined with an average maxRMSD >1.0Å, and a rigid cluster otherwise [20]). Same results were obtained using this classification (Figure 6B) (average lowest RMSD = 1.74Å and 1.52Å for flexible and rigid families, Wilcoxon p-value < 0.001).

### High flexibility regions are anti correlated with plDDT score

We mentioned that structural differences between conformers could be so tiny as the rotation of the side chains to large movements of loops and domains. This section studies how more flexible regions between apo and holo conformations are related to the plDDT score.

Using RMSF to measure protein flexibility between apo and holo conformers, we studied how this parameter correlates with plDDT. Taking only the models with the lowest RMSD to apo and holo forms, we found that the correlation between RMSF and plDDT is -0.39 (Pearson p-value < 0.001). However, this correlation increased when we used windows of different widths, reaching a maximum correlation of -0.45 (Pearson p-value < 0.001) with a window of 15 residues (Figure 7). RMSF captures the flexibility of the protein per position, as derived from the comparison between apo and holo conformers. To study how the intrinsic flexibility of each conformer relates with the plDDT, we used the profile of the normalized α carbons B-factors obtained by performing normal mode analysis for the apo form of the protein as described in Methods. In this way a similar correlation of -0.44 (Pearson p-value <0.001) was obtained.

**Figure 7:**
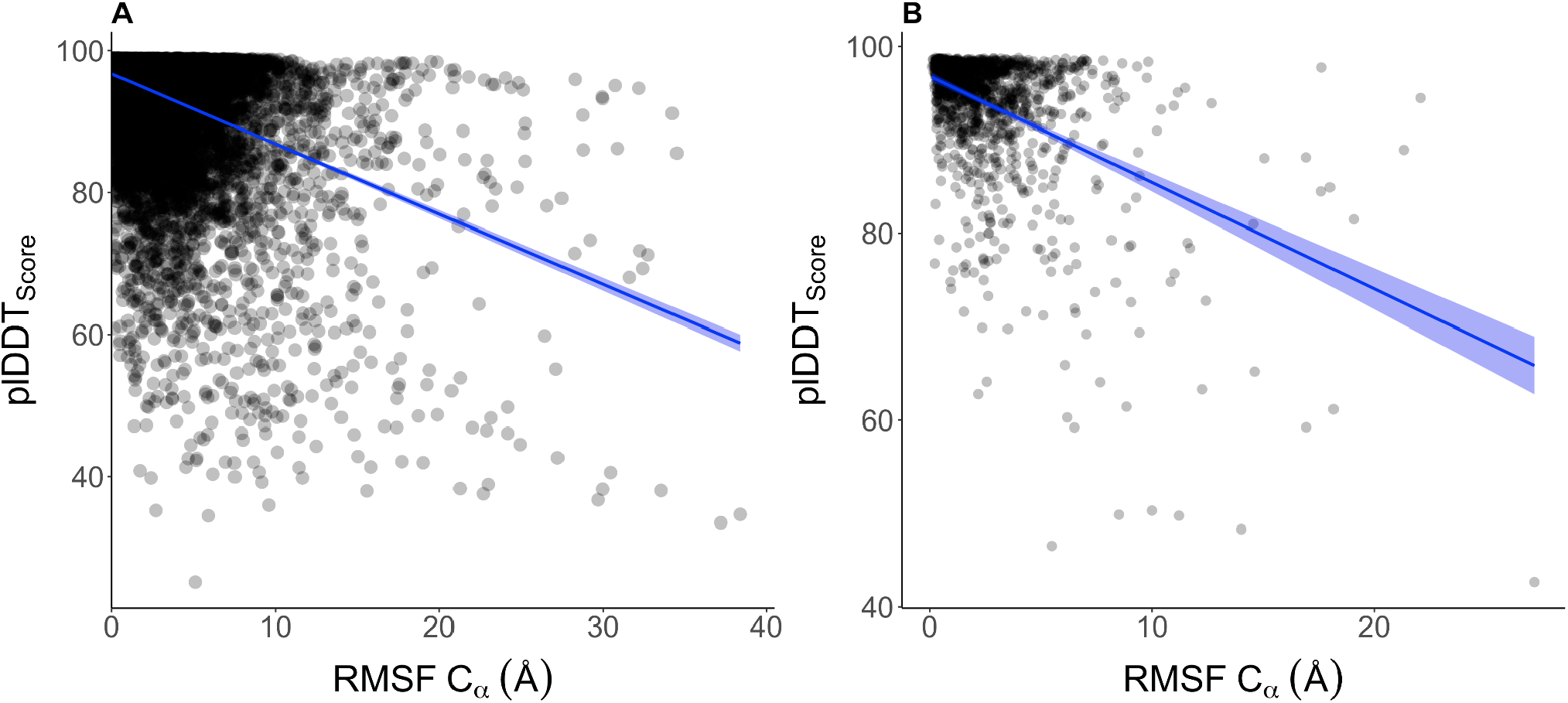
Distributions of plDDT score as a function of RMSF between apo and holo forms (left for all the positions and right using a window of 15 residues). Models were selected according to their lowest RMSD.

Low values of plDDT have been related to the occurrence of disordered regions [2]. However, according to our results, low-scoring regions could also represent flexible regions connecting ordered conformers, as observed for most of the proteins in our dataset.

## Discussion

AlphaFold’s breakthrough in predicting protein 3D models has certainly changed the way we study the protein structure-function relationship. Full structuromes of key organisms have been made available recently, along with easy-to-use utilities to run predictions. It is also outstanding to note that most of the predictions made are of the highest quality, mostly comparable with crystallographic resolution. In this work we have studied how the conformational diversity of the native state could be available through predictions and how in turn this key feature of protein biology affects the performance of predictions.

The first purpose is a practical one: Can we consider the top predicted models as snapshots of the conformational ensemble that describes the native state of proteins? Unfortunately, we can’t. In our dataset only 2% of the proteins showed models similar to either the holo or the apo forms with similar error, measured as the best RMSD to a given form (Figure 2). For the rest of the proteins, it is not possible to model both the apo and holo forms simultaneously with the same low error as when considering a single conformer (Figure 3). Far from disappointing, this observation was expected since a large set of redundant protein structures (conformers) would have been required during the neural network training processes [5] in order to predict conformational diversity.

We also found that most of the predictions made resemble and are mostly indistinguishable from the holo form of the studied protein (68% of the dataset). This is an exciting result because holo forms of proteins describe the binding capacity to a substrate or any other biologically relevant ligand (see Figure 4). Jumper et al. mentioned that AlphaFold could infer structures when the presence of a ligand is predictable from the sequence [2]. We thought that this finding could be explained due to a bias in the training process of AlphaFold2. However, exploring the BioLip database to estimate the relative presence of apo and holo forms in PDB shows about 64% of apo forms (a similar proportion observed in CoDNaS, from which our dataset was obtained). We hypothesized that holo forms could have a higher number of inter-residue contacts and that these could have influenced the modeling process in AlphaFold2. However, we did not detect differences in the number of contacts between holo and apo forms (the median number of contacts are equal to 3.47 and 3.44 of the apo and holo forms, Wilcoxon p-value >0.5). Apparently, differences in the number of directional polar interactions in contrast to interactions between nonpolar residues can explain differential flexibility patterns between holo and apo forms [12,38]. At this point, further work is required to understand this bias fully.

Does conformational diversity affect AlphaFold predictions? This second purpose of our work was a conceptual one related to the capability to recover evolutionary information related to protein flexibility encoded in multiple alignments. We have found that AlphaFold prediction capacity worsens with increasing conformational diversity of the protein being studied (Figure 5). We showed that this impairment is related to the heterogeneous dynamic behaviours in homologous protein families. Additionally, proteins from flexible homologous families are also difficult to predict (Figure 6). Several works showed that conformational diversity modulates the evolutionary process imprinting sequence information with dynamic behaviour [39–44]. Due to functional divergence, protein families could show different degrees of conformational diversity, making it difficult to extract specific sequence information from multiple sequence alignments for a given conformational motion. After the early and well-established observation that structures are very well conserved during evolution, it became evident that this conservation imposes structural constraints on sequence divergence [45–49]. However, and more recently, we showed that this sequence-structure relationship becomes fuzzy within families with significant degrees of conformational diversity [36]. Moreover, in protein families with complex dynamical behaviour (i.e., different degrees of conformational diversity), coevolutionary analysis allowed to infer inter residue contacts representing the most populated contacts among the family’s different structures, challenging the extraction of sequence features characterizing specific conformational patterns [37]. It is then expected that proteins belonging to families with heterogeneous flexibility behaviour would be difficult to predict from the evolutionary information used by AlphaFold2 (Figure 6 A and B).

Finally, our results suggest that the plDDT score can be used to scan flexible regions between ordered conformers. It was pointed out that plDDT could be helpful to predict disordered regions, but we can speculate that, as there is a continuum in ordered-disordered proteins [50], there could exist a range of plDDT thresholds to detect different sorts of protein flexibility. All the proteins in this work are mostly ordered (less than 15% of disordered regions), with regions with different flexibility. A ∼0.45 correlation between RMSF and plDDT in Figure 7 indicates that plDDT could capture the presence of flexible regions defining the conformational plasticity between apo and holo forms.

We think that our results provide tentative guidelines for practical uses in predicting 3D models of proteins using AlphaFold2 and adds a novel approach to a growing corpus of publications that nourishes the field by improving this extraordinary tool.

## Materials and Methods

### Description of the dataset

The set of apo and holo structures was obtained from the database of Conformational Diversity in the Native State of proteins (CoDNaS) [28]. CoDNaS is a redundant collection of PDB structures for the same sequence that can be taken as snapshots of protein dynamism. The conformational diversity for each protein was estimated as the C*α*-RMSD between apo and holo forms. In order to obtain a well-curated dataset containing protein motions related to a given biological activity we followed several specific quality criteria: (i) Only crystal structures with resolution < 3.9Å were considered; (ii) structures must not have missing residues; (iii) there must be 100% sequence identity between the conformers; (iv) structural deformations between pairs of conformers were associated with a given biological process based on experimental evidence; (v) no reported mutations; (vi) less than 15% disordered regions according to MobiDB consensus; and (vii) visual inspection was used to confirm an existing conformational diversity (e.g., movements should not be limited to flexible ends or arise from errors in the structural alignment). This allowed us to finally obtain apo-holo pairs of conformers for a total of 86 protein structures.

### Predictions and comparison of structures

Predicted models for each protein in the dataset were obtained using ColabFold ([6]) due to its easy access through Google Colab Notebooks. Runs were performed using no templates, automatic alignments and Amber energy minimization. For each run we used the 5 top models derived from the energy minimization. Each model was structurally compared between each other and against the correspondent apo and holo structures. As sequences between conformers and models are identical, the alignments are straightforward. We then quantified the structural similarity using the C*α* Root Mean Square Deviation (C*α*-RMSD).

### Evolutionary information

We sequentially clustered the CoDNaS database into homologous families containing sequences with more than 40% sequence identity and 70% coverage using CD-HIT. Each protein in CoDNaS has an associated maximum RMSD (maxRMSD) derived from the pairwise comparison of all its conformers. The maxRMSD is taken as the extent of the protein conformational diversity. A total of 175 well-populated clusters were taken (>8 proteins per cluster). A random protein from each cluster was modeled using ColabFold following the procedure mentioned above. The error of this model estimation was calculated as the lowest RMSD obtained from the comparison of any of the top 5 models with the crystallographic structures of the protein.

### B-factors analysis

Temperature factors or B-factors (*B*_*i*_) have been obtained performing normal mode analysis (NMA) using the coarse-grained Elastic Network Model [51,52] that considers the protein as an elastic network with nodes linked by springs within a cutoff distance r_c_. Herein the Cα are taken as nodes, and the value of r_c_ is varied from 7Å to 15Å for X-ray structures in order to optimize the correlation between theoretical and experimental B-factors, while r_c_=11Å is used for NMR structures. We perform the NMA for the apo form of the protein on the basis that normal modes obtained with the apo form of a given protein give a better description of the conformational change than those obtained with the holo form [53]. The normalized B-factor *B*_*i*_^*’*^ of atom *i* is obtained as *B*_*i*_ ^*‘*^ =(*B*_*i*_ -<*B*>)/σ(*B*), being <*B*> and σ(*B*) the average and standard deviation of the B-factor distribution for the corresponding protein structure, respectively. Each *B*_*i*_ ^*‘*^ was averaged over the neighbors of the ith residue within a radius of 7Å.

### Inter residue contacts analysis

Inter residue contacts have been obtained using the RING 2.0 web server [54]. Interacting pairs were identified following the closest contact strategy, i.e., all atoms are included to measure distances between residue pairs. While every pair of residues forms multiple interactions, the most energetic interaction per pair was considered. Interactions were defined distinguishing disulfides, salt bridges, hydrogen bonds, and aromatic interactions from generic van-der-Waals contacts.

### Motions classification

We have used the DynDom software v1.5 [35] to classify our dataset into proteins with “domain movements” (two or more domains presenting hinge movements) and proteins with “loop movements” (one domain, movements due to loops).

### Apo-holo characterizations

Classification of the 86 proteins in the dataset was done by manual curation following the bibliography. In parallel, the database of biological ligands BioLip [55] in its most recent version (October 01, 2021) was used to crosslink all the chains of the PDB (October 2021, total chains 661494) and CoDNaS v3 (March 2021, total chains 430151). If at least one biological ligand is found for a chain, it is assigned in the holo category; otherwise, it is considered as an apo conformer.

## Acknowledgements

NP, MSF, SFA, and GP are researchers, TS, JM, AJVR are postdoctoral fellows, and NE and EG are doctoral fellows from CONICET. This work was supported by Universidad Nacional de Quilmes (PUNQ 1309/19), ANPCyT (PICT-2018-3457), and the European Union’s Horizon 2020 Research and Innovation Programme (Grant Agreement No 778247). The funders had no role in study design, data collection and analysis, decision to publish, or preparation of the manuscript.

## Supplementary Figures and tables

Supplementary Table 1: [PDBs used in the dataset](https://drive.google.com/file/d/1GR-9cS5qYiJ9Ws7zJJcNe2nKf2ftbby9/view)

**Supplementary Figure 1:**
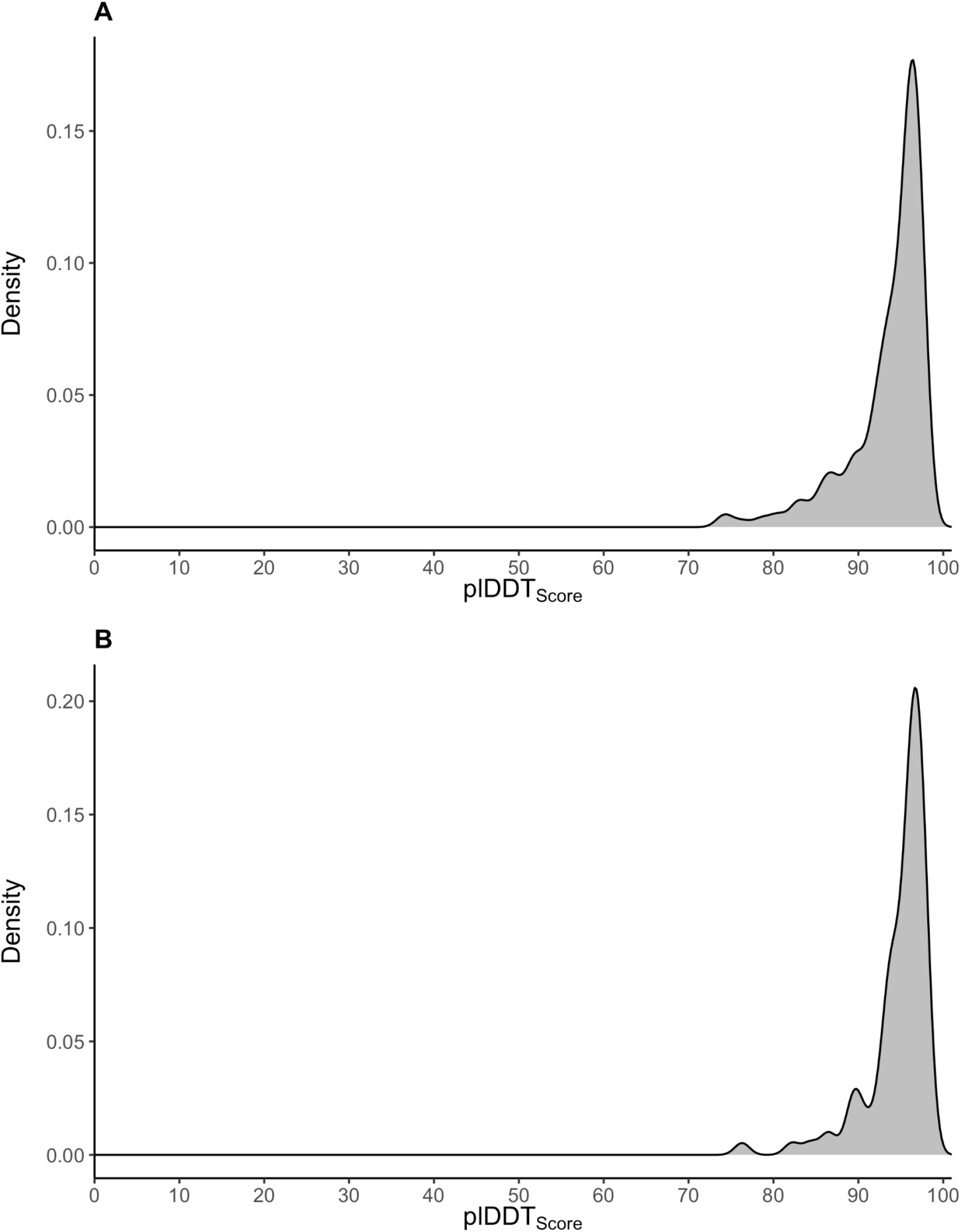
Distribution of plDDT scores for all the models obtained (**panel A**) and for the best model obtained (**panel B**).

**Supplementary Figure 2:**
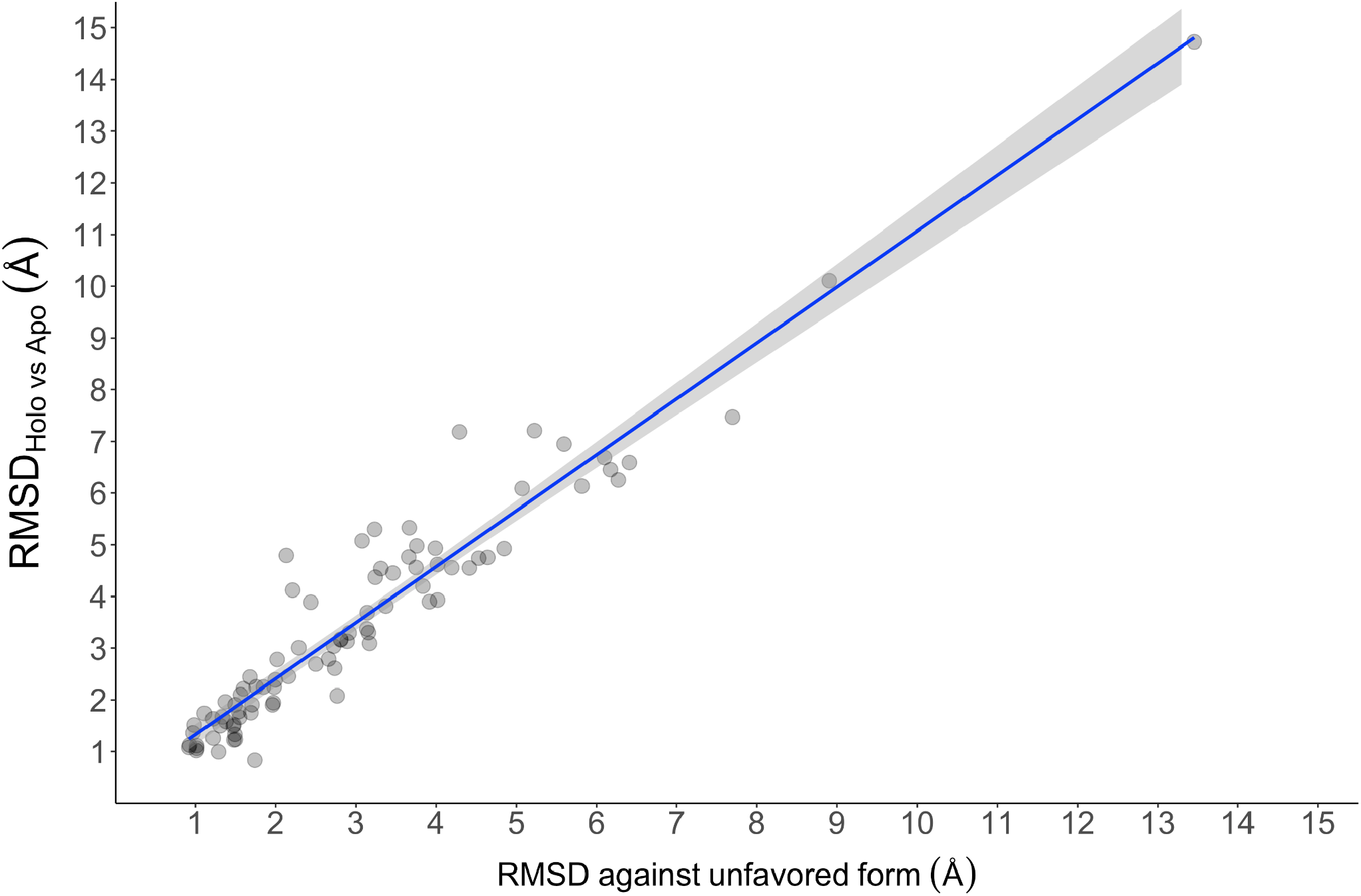
Correlation between the conformational diversity of the protein with the best RMSD measured to the unfavored form.

